# Cryo-EM reveals the dynamic interplay between mitochondrial Hsp90 and SdhB folding intermediates

**DOI:** 10.1101/2020.10.06.327627

**Authors:** Yanxin Liu, Daniel Elnatan, Ming Sun, Alexander G. Myasnikov, David A. Agard

## Abstract

TRAP1 is a mitochondrion specific Hsp90, a ubiquitous chaperone family that mediates the folding and maturation of hundreds of “client” proteins. Through the interaction with client proteins, TRAP1 regulates mitochondrial protein homeostasis, oxidative phosphorylation/glycolysis balance, and plays a critical role in mitochondrial dynamics and disease. However, the molecular mechanism of client protein recognition and remodeling by TRAP1 remains elusive. Here we established the succinate dehydrogenase B subunit (SdhB) from mitochondrial complex II as a client protein for TRAP1 amenable to detailed biochemical and structural investigation. SdhB accelerates the rate of TRAP1 dimer closure and ATP hydrolysis by 5-fold. Cryo-EM structures of the TRAP1:SdhB complex show TRAP1 stabilizes SdhB folding intermediates by trapping an SdhB segment in the TRAP1 lumen. Unexpectedly, client protein binding induces an asymmetric to symmetric transition in the TRAP1 closed state. Our results highlight a client binding mechanism conserved throughout Hsp90s that transcends the need for cochaperones and provide molecular insights into how TRAP1 modulates protein folding within mitochondria. Our structures also suggest a potential role for TRAP1 in Fe-S cluster biogenesis and mitochondrial protein import and will guide small molecule development for therapeutic intervention in specific TRAP1 client interactions.

## Introduction

Heat Shock Protein 90 (Hsp90) is a highly conserved molecular chaperone^1,2^. Coupled with ATP binding and hydrolysis, the Hsp90 dimer undergoes a conformational cycle between open and close states that facilitates the folding and maturation of hundreds of client proteins^3^. These clients are highly enriched in signal transduction and regulatory proteins, such as protein kinases, transcription factors, and E3 ligases^2,4^. In higher eukaryotes there are four Hsp90 homologs: the constitutively expressed cytosolic Hsp90β, the stress induced cytosolic Hsp90α, the endoplasmic reticulum localized Grp94, and the mitochondria localized TRAP1. The molecular mechanism of how Hsp90 mediates client protein maturation is poorly understood^5,6^. In the cytosol, the process is assisted by more than 20 cochaperones that modulate client binding and the ATP cycle^7^. By contrast, no cochaperones have been identified for the mitochondrial TRAP1. How TRAP1 can bypass the need for cochaperones to recognize and remodel its unique set of client proteins remains elusive.

One of the few identified *bona fide* client proteins for TRAP1 is Succinate Dehydrogenase B (SdhB)^8,9^, the B subunit of the mitochondrial respiratory complex II. SdhB incorporates three iron-sulfur (Fe-S) clusters and is a critical component of the mitochondrial electron transfer chain^10^. Here we biochemically reconstituted a TRAP1:client protein interaction system by using a truncated version of SdhB. The truncated SdhB accelerates both TRAP1 dimer closure and ATP hydrolysis rates. Cryo-EM structures of TRAP1 with and without SdhB bound revealed that SdhB goes through the lumen of a closed TRAP1 dimer and induces a conversion of the asymmetric closed state to near perfect symmetry^11,12^. This structure reveals a highly conserved Hsp90:client protein interaction mode that transcends the need for co-chaperones^6^. TRAP1 in turn stabilized several SdhB folding intermediates that are potentially beneficial for SdhB folding, Fe-S cluster biogenesis, mitochondrial complex II assembly, and protein import into mitochondria.

### Truncated SdhB accelerates TRAP1 closure and ATP hydrolysis

Our initial attempts to express and purify full-length recombinant human SdhB from *E. coli* suffered from severe degradation (Fig S1A), likely due to insufficient chaperone machinery to facilitate SdhB folding and Fe-S cluster loading^13,14^. Interestingly, numerous degradation products were capable of accelerating the TRAP1 closure rate as monitored using an inter-protomer FRET assay (Fig. S1)^12,15^. Client accelerated Hsp90 closure has not been observed with eukaryotic cytosolic Hsp90s but is indicative of productive client engagement for HtpG (bacterial Hsp90)^16,17^. Our FRET results suggested that truncated versions of SdhB can directly and efficiently interact with TRAP1, providing a potential model system for client recognition. Mass spectrometry of the SdhB degradation products determined that significant cleavage occurred near K160, L157, and F146. We chose to work on the K160 truncation which represented the dominant cleavage site and the longest construct among the three. As shown in Fig. 1A, this truncated construct of SdhB includes the complete N-terminal domain (NTD, residues 29-142), helix II (α2, residues 143-154) of the C-terminal domain, and a short loop region (residues 155-160).

**Figure 1.**
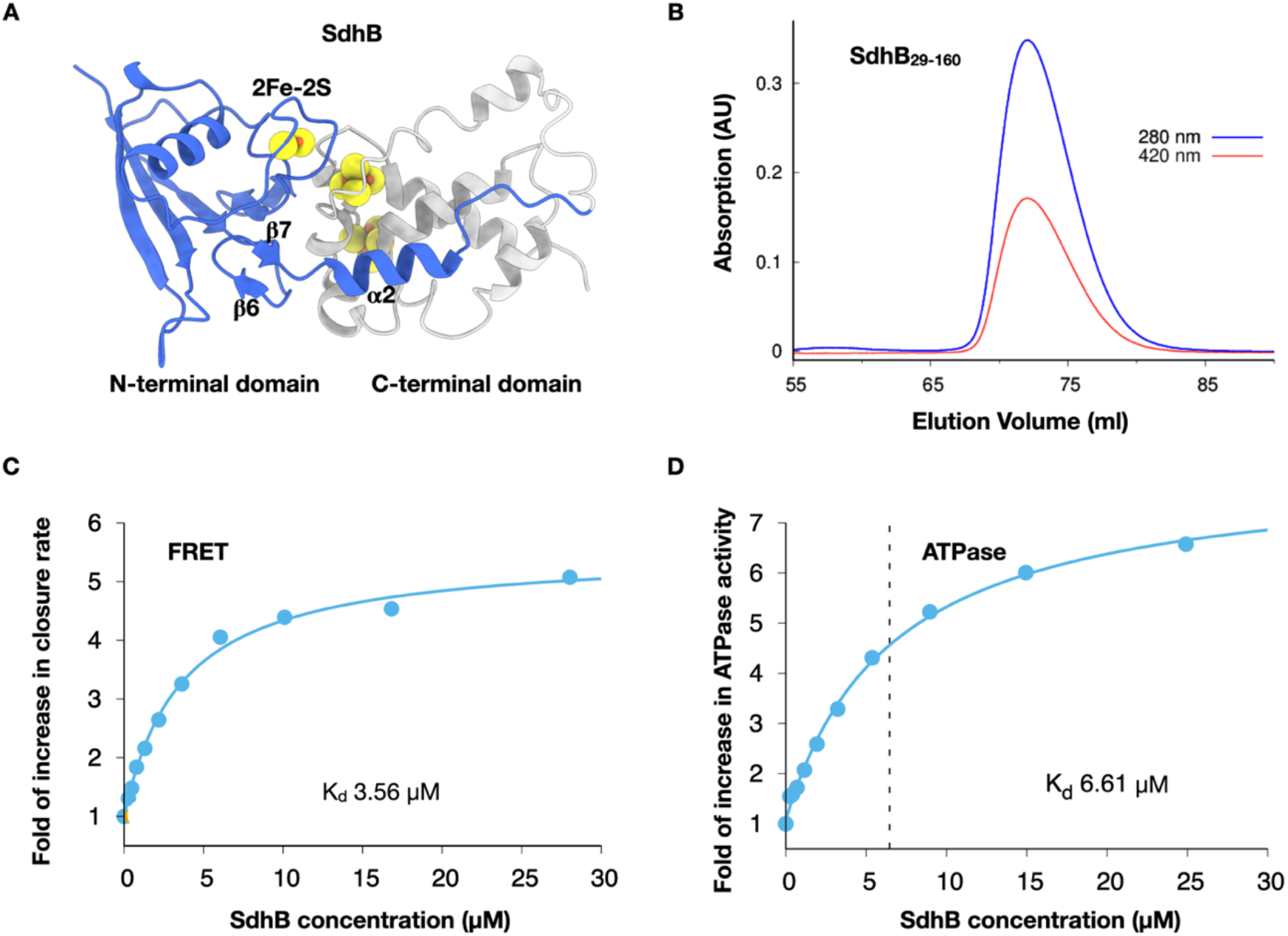
Establishment of SdhB_160_ as a biochemically tractable client protein for TRAP1. (A) The native state of SdhB from previous crystal structure of mitochondrial complex II. Three Iron-sulfur clusters are highlighted in sphere representation. The truncation version SdhB_160_ used in current work is colored blue. (B) Gel filtration of SdhB_160_ construct show absorbance at 420 nm, indicating the binding of 2Fe-2S cluster. (C) SdhB_160_ accelerate the closure of TRAP1 dimer in FRET assay. (D) SdhB_160_ accelerate the ATP hydrolysis in TRAP1.

The corresponding truncated construct (SdhB_160_) expressed well and was readily purified to homogeneity without signs of degradation (Fig. S1D). SdhB_160_ was brown in color with an absorption peak around 420 nm (Fig. 1B), consistent with binding of the one 2Fe-2S cluster located in the NTD (Fig. 1A)^10^. Similar to the results obtained with the mixture of degradation products, SdhB_160_ accelerated TRAP1 closure by 5-fold under saturating conditions with an EC_50_ of 3.6 μM (Fig. 1C). Since dimer closure is the rate limiting step for TRAP1 ATP hydrolysis^15^, SdhB_160_ should also accelerate TRAP1 ATPase activity, which it does with an EC_50_ of 6.6 μM (Fig 1D). It was observed previously that HtpG closure and ATPase were accelerated by model client proteins such as Δ131Δ staphylococcal nuclease^16^ or ribosomal protein L2^17^. Thus, we here established SdhB as a *bona fide* client protein having an equivalent activating effect on TRAP1.

### SdhB induces an asymmetric to symmetric transition for TRAP1

To better understand the interplay between TRAP1 and SdhB, we determined cryo-EM structures for the complex. Our FRET experiments indicated that SdhB stimulates formation of the closed state of TRAP1, suggesting that it would also bind that state. Therefore, we used the non-hydrolyzable ATP analog AMP⋅PNP to stabilize the TRAP1:SdhB_160_ complex in the closed state. Image analysis (Fig. S2) revealed two structures representing the closed TRAP1 dimer with and without SdhB bound, which were further refined to 3.14 Å and 3.26 Å, respectively (Fig. 2 and Fig. S3). Previously determined crystal structures of zebrafish TRAP1 with various ATP analogs and a cryo-EM structure of human TRAP1 with ADP⋅BeF_x_ revealed an asymmetric closed state for TRAP1^11,18^. Here the client-free AMP⋅PNP bound human TRAP1 adopted the same asymmetric state with one protomer in straight conformation and the other protomer in a characteristic buckled conformation (Fig. 2A).

**Figure 2.**
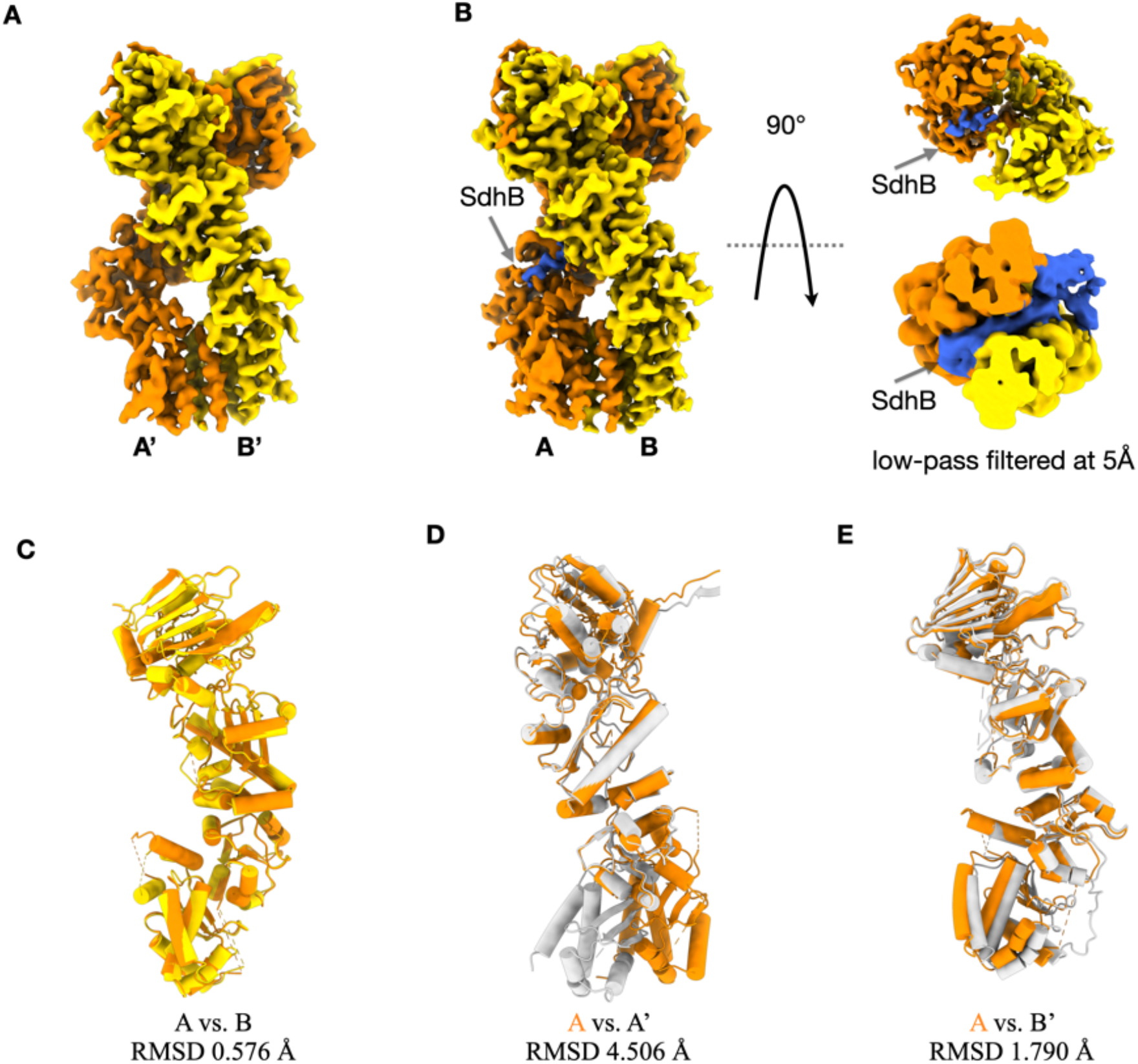
Cryo-EM structures show an asymmetric to symmetric transition for TRAP1 induced by SdhB binding. (A) Cryo-EM structure of human TRAP1 closed state. (B) Cryo-EM structure of human TRAP1 in complex with client protein SdhB. (C) Alignment between two protomers of TRAP1 in the SdhB bound states. (D) Alignment between the bulked protomer of TRAP1 in the asymmetric closed state (A’ in (A), gray) and the protomer A of TRAP1 in the SdhB bound symmetric state (A in (B), orange). (E) Alignment between the straight protomer of TRAP1 in the asymmetric closed state (B’ in (A), gray) and the protomer A of TRAP1 in the SdhB bound symmetric state (A in (B), orange).

The second structure represented the TRAP1:SdhB_160_ complex (Fig. 2B). When low pass filtered to 5 Å, the SdhB density became clearly visible in the lumen region and on one side of the TRAP1 dimer. Surprisingly, upon binding to SdhB, TRAP1 was converted to a symmetric closed state that had not been previously observed. The RMSD between the two protomers in the client binding state is only 0.58 Å (Fig. 2C). The symmetric protomers are dramatically different from the buckled protomer in the asymmetric state, having an RMSD of 4.5 Å as shown in Fig. 2D. While both protomers more closely resemble the straight arm from the asymmetric state (RMSD of 1.8 Å), nonetheless there are noticeable differences in the rotation angle between the TRAP1 middle domain and the C-terminal domain, corresponding to a further twisting of the CTD (Fig. 2E). In summary, client protein binding induced a completely unexpected asymmetric to symmetric structural conversion in TRAP1. A similar symmetric Hsp90 state was observed when cytosolic Hsp90s bind cochaperones (such as p23^19^ or Aha1^20^) or client proteins (such as the Hsp90:Cdc37:Cdk4 kinase^6^ or Hsp90:p23:Glucocorticoid receptor complexes^21^). Together they suggest a universal utilization of a symmetric Hsp90 closed state for the final stages of client binding regardless of the presence cochaperones.

### Partly unfolded SdhB_160_ goes through the lumen of TRAP1

Although the overall resolution for the TRAP1:SdhB_160_ complex reached 3.14 Å, the resolution of the SdhB region was limited. Consequently, we carried out focused classification^22^ on the globular region of the SdhB_160_ and obtained four distinct structures (Fig. 3 and Fig. S4). The TRAP1 regions were well resolved and the lumenal SdhB_160_ densities were clearly visible in all four structures. However, the conformations of the client globular regions were quite different from one another, highlighting the conformational heterogeneity of SdhB_160_ when bound to TRAP1. We hypothesize that these structures represent various folding intermediate states of SdhB_160_ that were captured and stabilized by TRAP1.

**Figure 3.**
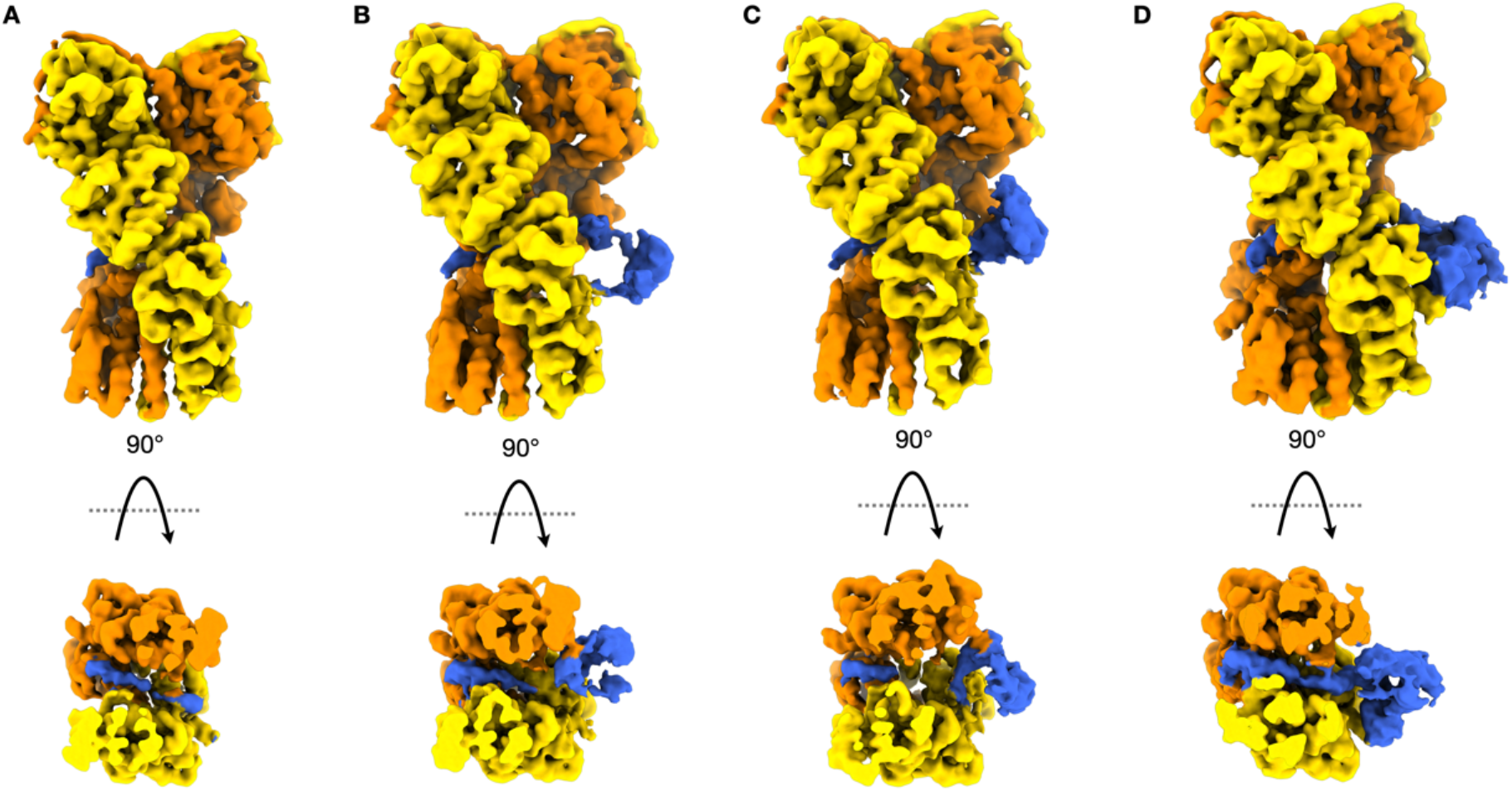
Cryo-EM structures of TRAP1 bound with SdhB in various folding intermediate states.

An atomic model of the TRAP1:SdhB complex was built using the map shown in Fig. 3D, which had the best resolved SdhB region among the four structures. In this state, the SdhB-NTD was captured on one side of the TRAP1 as a globular density which is connected with a continuous density in the TRAP1 lumen (Fig. 3D and Fig. 4A,B). The resolution of the globular region remained limited to 5~6 Å despite extensive focused classification (Fig. S4D). However, the SdhB-NTD in the crystal structure can be docked in this region unambiguously (Fig 4A). The 2Fe-2S cluster stands out as the strongest density in the region as expected and is visible even at high contour level (Fig 4A inset). The SdhB-NTD interacts with TRAP1 with two contact surfaces. First, the M133 from SdhB is interacting with F621 from the TRAP1 amphipathic helix (Fig. 4C).

**Figure 4.**
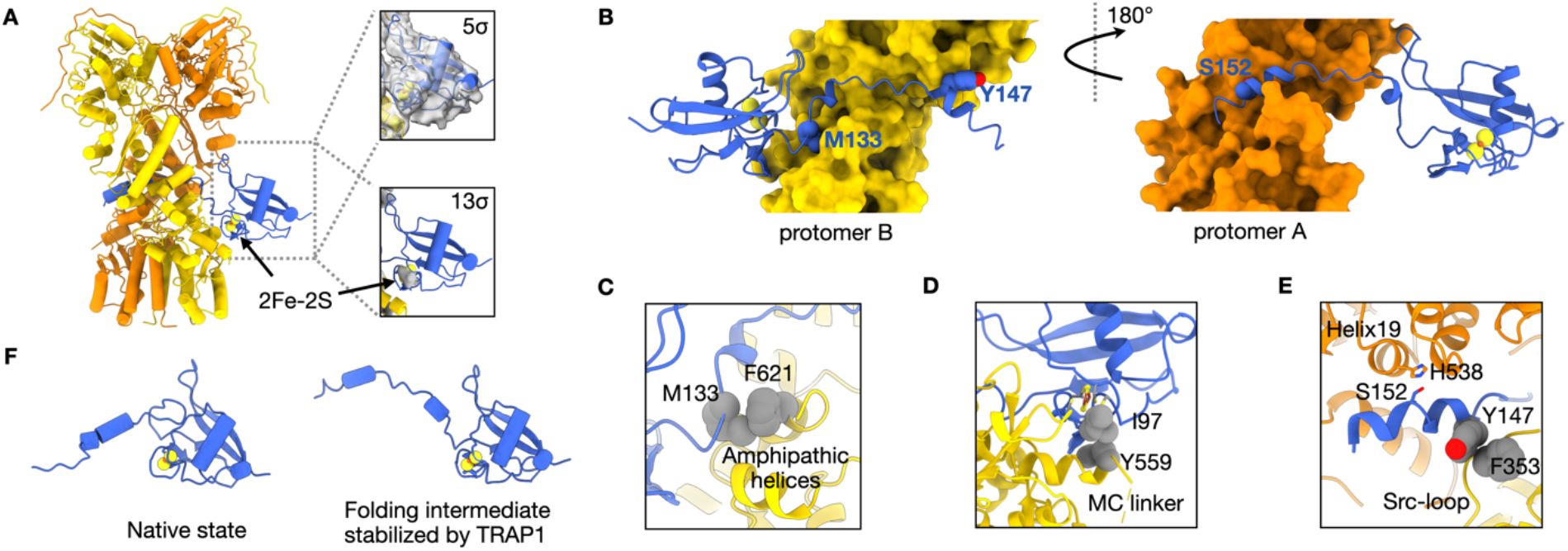
Interactions between TRAP1 and a partially folded SdhB. (A) Atomic model of TRAP1 in complex with a partially folded SdhB. The insets show the docking of native SdhB-NTD into the cryo-EM map at a low contour level 5σ and a high contour level 13σ, respectively. The strongest density corresponds to 2Fe-2S. (B) A stretch of SdhB-CTD goes through the TRAP1 lumen. (C) The first interaction site between TRAP1 (F621 in the amphipathic helix I) and SdhB (M133). (D) The second interaction site between TRAP1 (Y559 at the beginning of MC linker) and SdhB (I97). (E) The interactions between SdhB-α2 and TRAP1 involve both protomers, helix-19 from protomer A and Src-loop from protomer B. (F) The comparison between the native state of SdhB_160_ and the partially folded intermediate state stabilized by TRAP1.

The amphipathic helix in Hsp90 is known as a client interacting region^23^ and TRAP1^F621^ is highly conserved hydrophobic residue among the Hsp90 homologs (Fig. S5). Although F621 is replaced by a methionine in HtpG and human Hsp90s, it is known to interact with client proteins in both cases^6,23^. The second contact point is between SdhB^I97^ and TRAP1^Y559^ near the end of TRAP1 middle domain (Fig. 4D). This is a novel client interacting region that was not reported previously.

The resolution of the SdhB lumen density is sufficiently high to reveal a short helix in the lumen on the opposite side of the SdhB-NTD and the register can be unambiguously assigned. This helix corresponds to helix II (α2) in the SdhB-CTD (Fig. 1A and Fig. 4B). In fact, the SdhB-NTD alone (deletion of α2 and the following short loop) failed to accelerate TRAP1 closure (Fig. S6). The interactions between SdhB and TRAP1 within the lumen are predominantly hydrophobic (Fig. S7). Due to the narrow nature of the lumen (Fig. S8), SdhB asymmetrically interacts with both TRAP1 protomers. A hydrogen bond is formed between SdhB-α2^S152^ and TRAP1^H538^ from helix 19 in protomer A (Fig. 4E). The helix 19 from HtpG and human Hsp90b have been reported to interact with client proteins^6,23^. Hydrophobic interactions play a major role in stabilizing the interaction between SdhB-α2 and Src-loop from TRAP1 protomer A as shown in Fig. 4E. The Src-loop in cytosolic Hsp90s are known to be important for kinase maturation^6,24^. The observation that the TRAP1 Src-loop is directly involved in mitochondrial client maturation reveals the importance of this motif and thus represents a conserved underlying mechanism by which Hsp90s interact with their clients.

The atomic model (Fig 4A) shows that SdhB undergoes a partial unfolding (Fig. 4F). In order for SdhB-α2 to reach on the other side of TRAP1, a short β-harpin formed with β6 and β7 in the native structure of SdhB (shown in Fig. 1A) has to unfold and extend through the lumen of TRAP1. In fact, a 34-amino acid peptide derived from SdhB (SdhB_127-160_) is sufficient to accelerate the TRAP1 ATPase activity (Fig. S9), although not as efficiently as SdhB_160_. Thus, our model represents a nearly folded state of SdhB that is stabilized by TRAP1. All the other states shown in Fig 3A-C represent TRAP1-stabilized SdhB folding intermediates having various degrees of unfolding.

## Discussion

Together with previous cryo-EM structures of cytosolic Hsp90-client interactions (Hsp90:Cdc37:Cdk4 and Hsp90:p23:GR)^6,21^, our structures of the mitochondrial TRAP1:SdhB complex reveal broadly conserved modes of client binding that transcend stabilization by co-chaperones. In all three cases, the partially unfolded client proteins pass through the lumen of Hsp90. Notably, there is no requirement that the larger more folded domain of the client be N-terminal (SdhB) or C-terminal (Cdk4, GR), indicating that the lumen only requires a hydrophobic stretch of client. It is now clear that there are conserved client protein binding sites located in the Src-loop^23^, Helix 19, and the amphipathic helices from the Hsp90-CTD^6,21,23^. These amphipathic helices and adjacent loops emerge as highly adaptable binding surfaces that can present both hydrophobic and polar surfaces for client and cochaperone interactions. In contrast to its cytosolic counterpart, TRAP1 accomplishes its client protein maturation in the absence of a co-chaperones and undergoes a dramatic asymmetric to symmetric structural transition upon client binding. The buckled arm in asymmetric closed state of TRAP1 is incompatible SdhB binding (Fig. S10). This suggests that the asymmetric state may represent an on-pathway intermediate after the first ATP is hydrolyzed^12^, or possibly serve as a transient regulatory step in client protein remodeling and release.

There are important functional implications of our TRAP1:SdhB structures. First, by stabilizing SdhB folding intermediate states, TRAP1 will prevent the aggregation of partially folded SdhB formed either upon import or through transient unfolding events. Second, the observed binding mode would decouple the folding of the SdhB-NTD from the SdhB-CTD allowing each domain to fold independently on opposite sides of the TRAP1 dimer. Third, TRAP1 could serve as a scaffold protein to present a partially folded SdhB for iron-sulfur cluster loading and complex II assembly with the help of additional chaperones^14,25^. Finally, in binding to a newly translocated SdhB-NTD, TRAP1 may play a similar role as mitochondrial Hsp70 in promoting mitochondrial protein translocation^26^. In all these scenarios, TRAP1 would play an important role in mitochondrial protein homeostasis, metabolism, and the switch between oxidative phosphorylation and glycolysis^8,9,27,28^, through its direct interactions with SdhB.

## Acknowledgements

We thank members of the Agard Lab for helpful discussions. We gratefully thank David Bulkley, Eric Tse, Michael Braunfeld, and Glenn Gilbert from the W.M. Keck Foundation Advanced Microscopy Laboratory at the University of California, San Francisco (UCSF) for maintaining the EM facility and helping with data collection. Special thanks to Matt Harrington and Joshua Baker-LePain for computational support on the USCF Wynton cluster. This work was supported by funding from National Institutes of Health grants U54CA209891, S10OD020054, and S10OD021741. Y.L. was supported by a Howard Hughes Medical Institute-Helen Hay Whitney Foundation Postdoctoral Fellowship, an American Heart Association Postdoctoral Fellowship grant #18POST33990362, and the Program for Breakthrough Biomedical Research which is partially funded by the Sandler Foundation. M.S. was supported by 2018 AACR-Takeda Oncology Lymphoma Research Fellowship grant #18-40-38-SUN. D.A.A was supported by the Howard Hughes Medical Institute.

## Authors contributions

Y.L. and D.A.A. designed all the experiments. Y.L. and D.E. designed the construct, purified the proteins, and performed biochemical experiments. Y.L., M.S., and A.G.M. collected and processed the cryo-EM data. Y.L. built the atomic models. D.A.A. supervised the project. Y.L. and D.A.A wrote the manuscript and all authors contributed to editing.

## Data availability

The electron microscopy maps and atomic models have been deposited into the Electron Microscopy Data Bank (EMDB) and the Protein Data Bank (PDB). The accession codes are EMD-22811 and PDB-7KCK for TRAP1 structure without SdhB bound, EMD-22812 and PDB-7KCL for the consensus model of SdhB-bound TRAP1 structure. The accession codes for the maps of TRAP1 bound with SdhB in various folding intermediates are EMD-22813, EMD-22814, EMD-22815, and EMD-22816, respectively. The accession code for the atomic model derived from EMD-22816 is PDB-7KCM.

