## Supplemental Information for "Cryo-EM reveals the dynamic interplay between mitochondrial Hsp90 and SdhB folding intermediates"

This PDF file includes:

Methods
Figures S1 to S10
Tables S1
References

### METHODS

**Protein cloning, expression, and purification**

The human SdhB gene (residue 29-160) was codon optimized and cloned into pET28a expression plasmids with a N-terminal 6×His-tagged followed by a HRV 3C protease cleavage site. Expression was carried out in *E. coli* BL21(DE3)-RIL. Cells were grown in TB media (supplemented with 0.2 mM cysteine and 0.2 mM ferrous sulfate) at 37˚C to $OD_{600}$ of ~0.6 and then induced with 0.5 mM IPTG at 16 ˚C for 18 hr. Proteins were first purified by Nickel-affinity chromatography and incubated with HRV 3C protease overnight at 4˚C without any reducing agent to cleave the N-terminal His-tag. After confirming the His-tag cleavage by SDS-PAGE, proteins were further purified using Mono S cation exchange and size exclusion chromatography before they were aliquoted and snap frozen with liquid $N_2$. The construct, expression, and purification procedure for human Trap1 were as same as previously described[1,2].

**Steady-state ATPase assay**

The ATPase assay monitors phosphate release via a chromogenic substrate 7-methyl-6-thioguanosine (7-MESG) in the presence of an *E. coli* PNPase (purine nucleoside phosphorylase)[3]. In these assays, 0.14 mM 7-MESG and 2 μM PNPase was used per reaction. The change in absorbance of 7-MESG was measured at 355 nm. Initial rates from phosphate release assays were obtained from fitted slopes of a linear function, $y = mx + b$, to the most linear region of each kinetic trace. The activity *versus* concentration curves were analyzed by a standard Michaelis-Menten equation.

**Fluorescence resonance energy transfer (FRET) assay**

The FRET sample preparation and measurement was as same as described previously[2,4]. In short, Cysteines were introduced at Glu140 and Lys413 and labeled with Alexa Fluor 555 and Alexa Fluor 647 maleimide. For FRET experiments, 500 nM of labeled Trap1 were used in the presence of various concentration of wild type SdhB truncations. Fluorescence measurements were taken in the SpectraMax M5 plate reader with SoftMax Pro software for data acquisition. Samples were excited at 532 nm and emission wavelengths were collected at 567 nm and 668 nm for donor and acceptor fluorescence, respectively. Relative FRET efficiencies are calculated by taking the ratio of acceptor to donor fluorescence intensity.

**Cryo-EM sample preparation**

The closed state of Trap1 in complex with SdhB was obtained by incubating 2 μM Trap1 with 20 μM SdhB in the presence of 1 mM AMPPNP and 1 mM $MgCl_2$ at 30 °C for 30 min. Cryo-EM grids were prepared with Vitrobot Mark IV (FEI, Thermo Fisher Scientific, Hillsboro), using 16°C and 100% humidity. 4 μL aliquots of samples were applied to glow discharged Quantifoil R1.2/1.3, 300-mesh copper holey carbon grids (Quantifoil Micro Tools, GmbH, Großlöbichau, Germany), single blotted for 8 seconds with blot force 3, and plunge frozen in liquid ethane cooled by liquid nitrogen.

**Cryo-EM data collection**

Data were collected on a Titan Krios microscope (Thermo Fisher Scientific) operated at 300 kV with a K2 Summit direct electron detector (Gatan, Inc.). A GIF-BioQuantum energy filter with a slit width of 20 eV was used. Images were recorded using SerialEM[5] with defocus varied from -0.5 to -2.1 μm for a total dose of 72 $e^-/Å^2$. A super-resolution pixel size of 0.407 Å was used and each image was dose-fractionated to 100 frames (0.1 s each, total exposure of 10 s) with a dose rate of 7.2 $e^{-}/Å^{2}/s$. A summary of the data collection parameter was provided in Supplementary Table 1.

**Image processing**

Image stacks were motion-corrected and summed using MotionCor2[6], resulting in Fourier-cropped summed images which are binned by 2. CTFFIND4 was used to estimate defocus parameters for all the images[7]. Initial particle picking was carried out using Gautomatch without a template to generate the 2D class averages, which were then used as templates for a second-round particle picking on micrographs with 25 Å low-pass filtering. Relion 3.1[8,9] was used for all the following steps. One round of reference-free 2D classification were performed for 25 iterations each with images binned by 4. Good particles were picked from 2D averages, extracted as images binned by 2, and subjected to 3D classification within Relion. An initial model was generated from cryo-EM map of human Trap1 (EMD-22174 and PDB 6XG6) and low-pass filtered to 30 Å. Classes corresponding to Trap1 alone and Trap1:SdhB complex were re-extracted without binning, and subjected to 3D auto-refinement. Focused classification with signal subtraction were carried out for the Trap1:SdhB complex with a spherical mask with 25 Å diameter around the SdhB region. All refined maps were post-processed and sharpened by an automatically estimated B-factor. All resolutions were estimated by applying a soft mask around the protein density and the gold-standard Fourier shell correlation (FSC) = 0.143 criterion.

**Model building, refinement and validation**

The initial model of Trap1 was derived from the cryo-EM structure of ADP·$BeF_x$ bound human Trap1 (PDB code 6XG6). The nucleotide was replaced with AMP·PNP. The initial model of human SdhB were derived from crystal structure porcine SdhB (PDB code 1ZOY). The initial models of Trap1:SdhB complexes were generated by rigid-body-docking individual domains of Trap1 and SdhB-NTD into the cryo-EM maps using UCSF ChimeraX[10]. The docked models were refined in real space against the cryo-EM maps using real space refinement in PHENIX[11] with secondary structure restraints, followed by iterative rounds of manual and automated refinement in Coot[12] and PHENIX[11], respectively. The final models and their fitness into the maps were visually inspected and geometry was further evaluated using MolProbity[13]. Cryo-EM data collection, refinement and validation statistics are summarized in Supplementary Table 1. Figures depicting the structures were prepared in ChimeraX[10] or VMD[14].

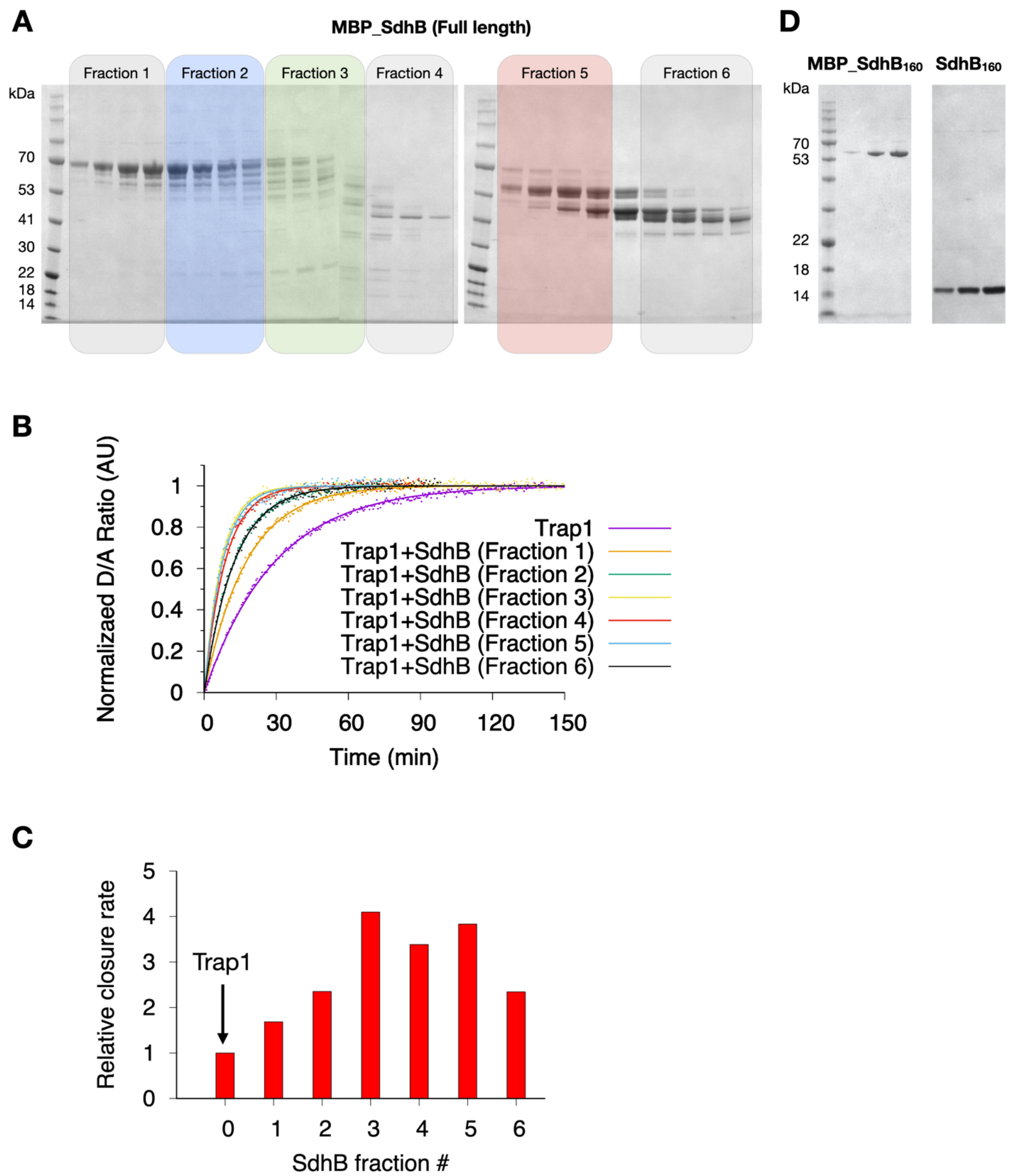


**Figure S1.** SdhB degradation products accelerate Trap1 closure rate. (A) SDS-PAGE shows severe degradation of full-length SdhB after purification through anion exchange and size exclusion chromatography. The degradation products were collected into six fractions. (B) All fractions of degradation products accelerate Trap1 closure rate as shown in FRET assay. (C). Quantification of the Trap1 relative closure rate measured in FRET. (D) Purified $SdhB_{160}$ with and without MBP tag shown in SDS-PAGE. The latter was used in this study.

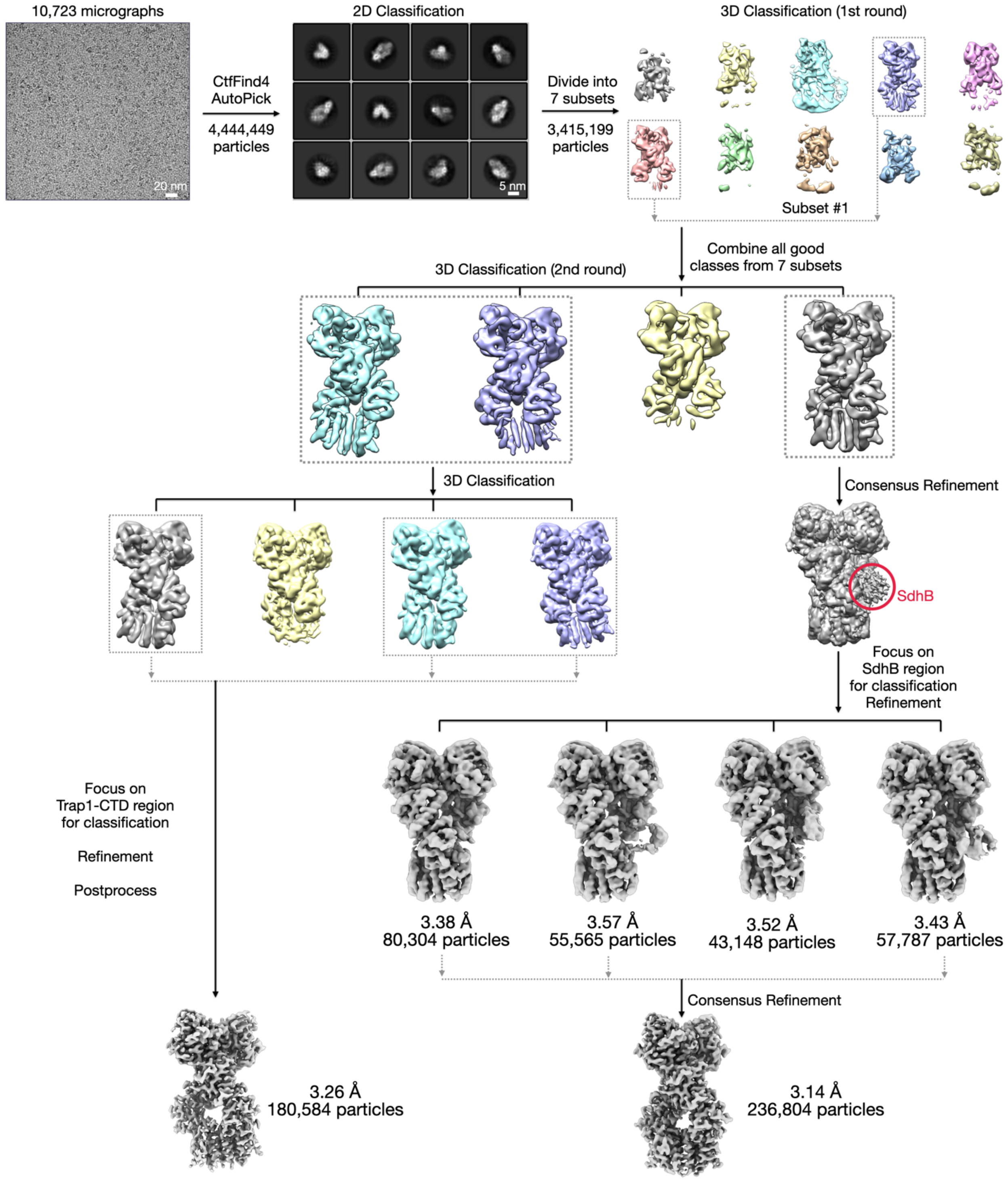


**Figure S2.** Cryo-EM workflow for image analysis.

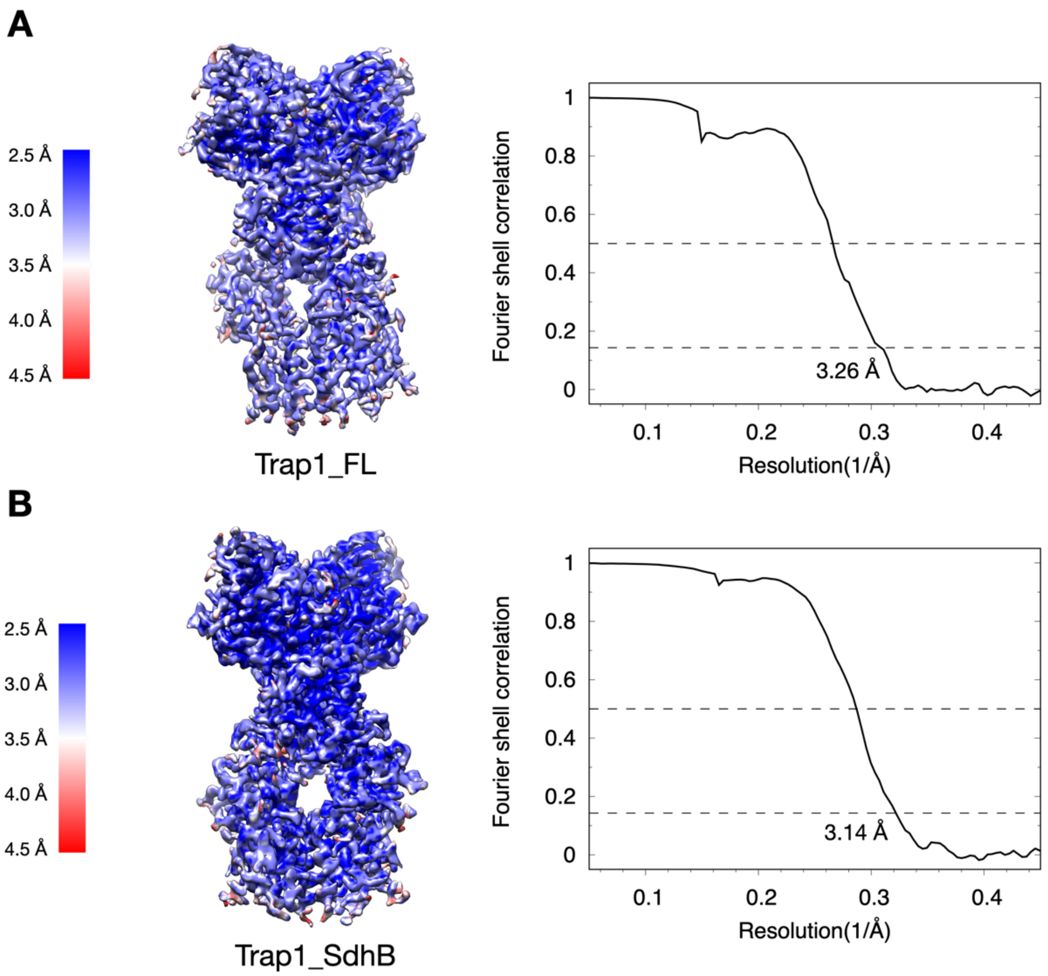


**Figure S3.** Local resolution map and gold-standard Fourier shell correlation (FSC) curves for cryo-EM structures of Trap1 (A) without SdhB bound and (B) with SdhB bound.

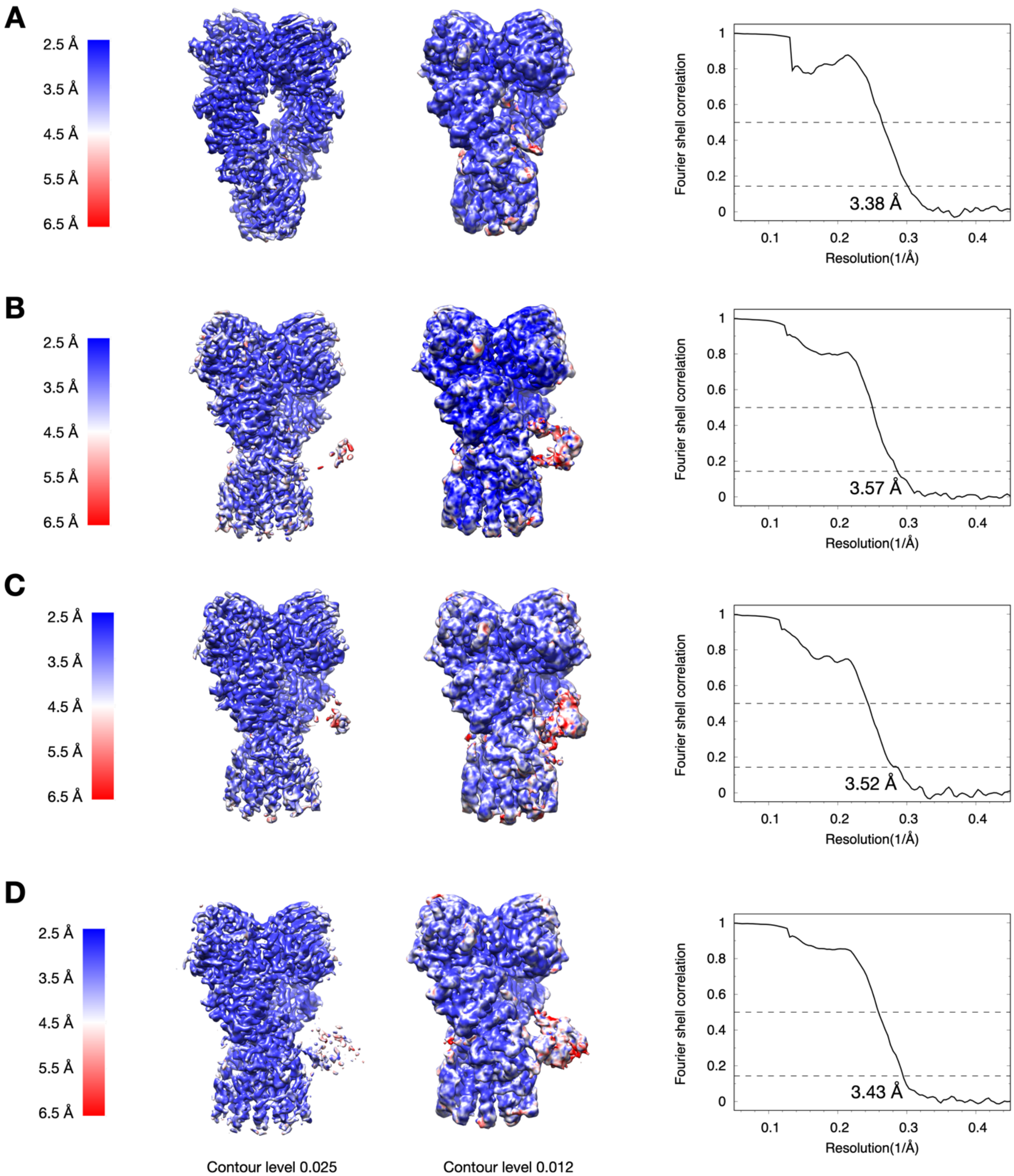


**Figure S4.** Local resolution map at the different contour levels and gold-standard Fourier shell correlation (FSC) curves for four cryo-EM structures of SdhB-bound Trap1 resolved through focused classification.

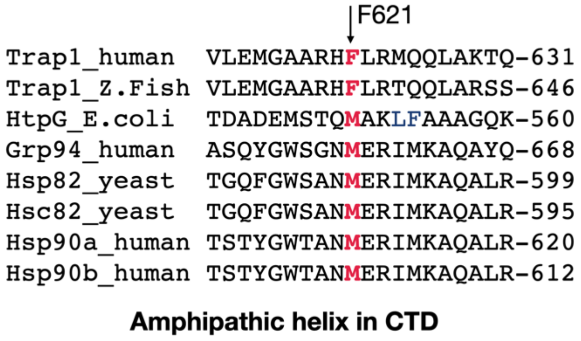


**Figure S5.** Sequence conservation of amphipathic helix in CTD among all Hsp90 homologs. F621 is highly conserved client protein interacting residue.

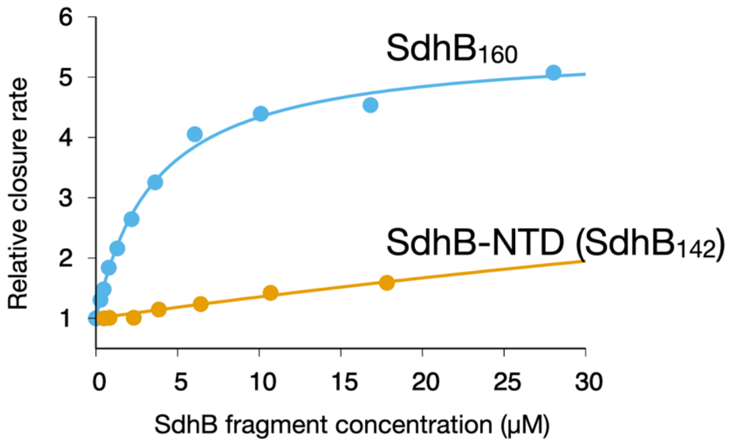


**Figure S6.** Truncation of α2 and the short loop afterward in $SdhB_{160}$, resulting in the construct of SdhB-NTD ($SdhB_{142}$), abolished its accelerating effect on Trap1 closure measured by FRET assay.

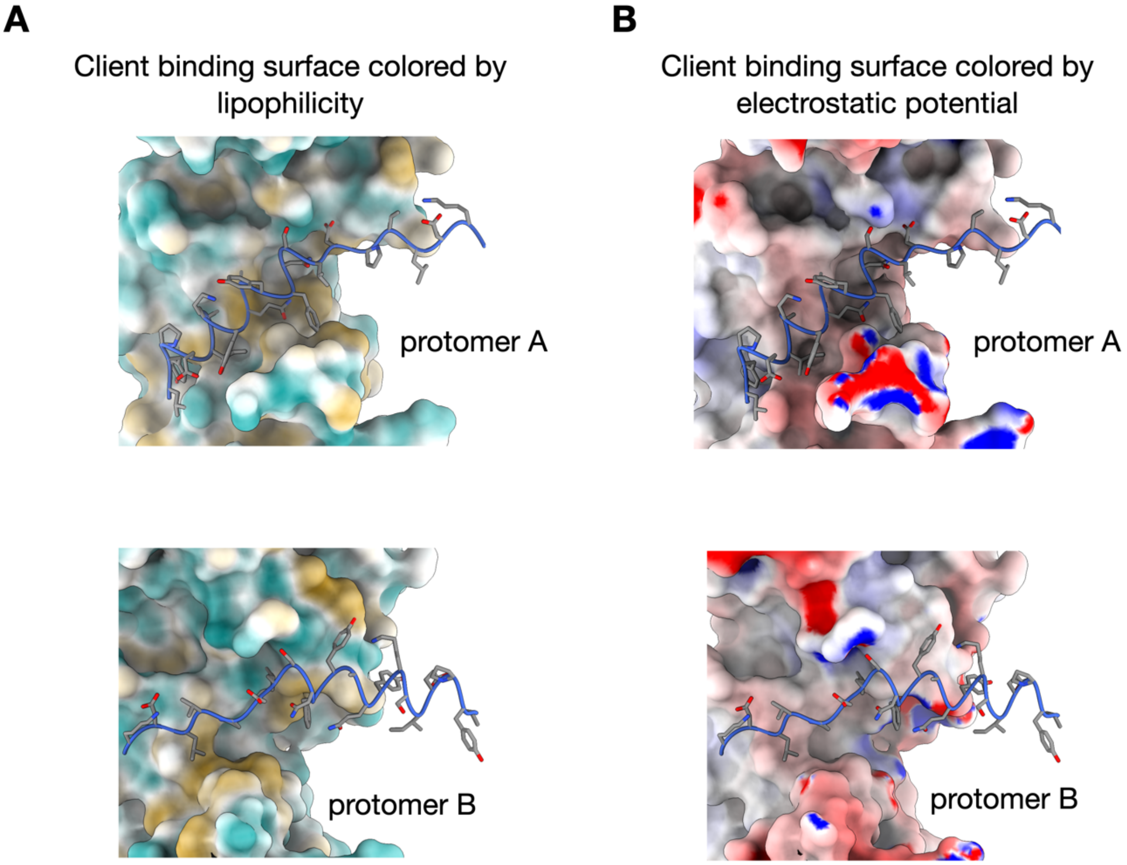


**Figure S7.** Client binding surface in the Trap1 lumen colored by (A) lipophilicity and (B) electrostatic potential. Cyan and golden represent hydrophilic and hydrophobic regions, respectively. Red and blue represent negatively and positively charged regions, respectively. The SdhB is shown in cartoon and stick style.

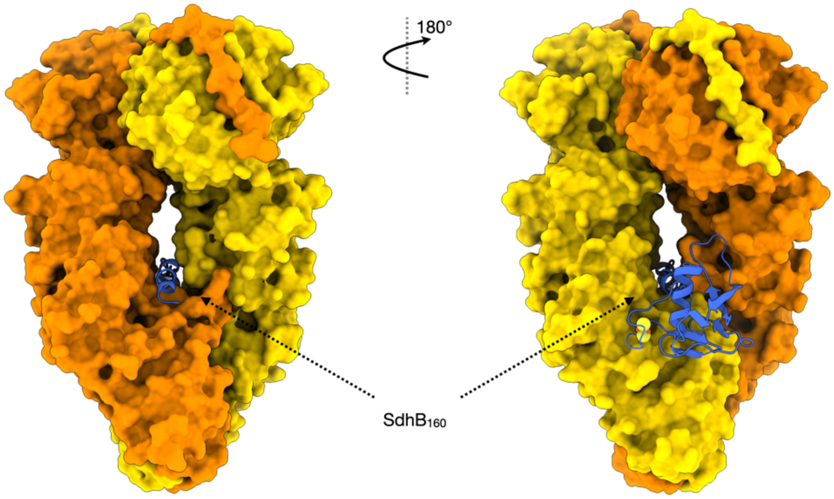


**Figure S8.** The sideview of TRAP1:SdhB$_{160}$ complex shows the SdhB$_{160}$ goes through the narrow lumen of TRAP1.

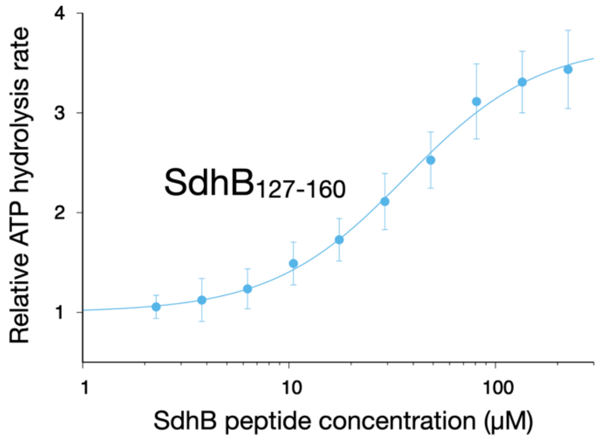


**Figure S9.** The lumen binding 34-amino acid peptide from SdhB ($SdhB_{127\text{-}160}$) accelerates the ATPase activity of Trap1.

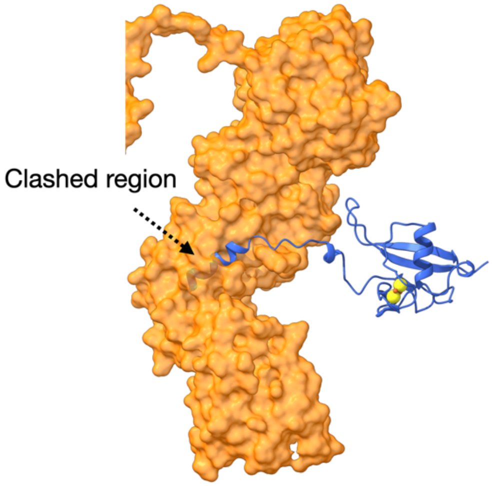


**Figure S10.** The asymmetric TRAP1 closed state is incompatible with $SdhB_{160}$ binding. After aligning on the straight protomer B (not shown), a clash is observed between the buckled protomer A and $SdhB_{160}$.

### **Table S1.** Cryo-EM data collection, refinement and validation statistics

| | #1 Trap1 (EMD-22811) (PDB 7KCK) | #2 Trap1_SdhB$_0$ (EMD-22812) (PDB 7KCL) | #3 Trap1_SdhB$_1$ (EMD-22813) | #4 Trap1_SdhB$_2$ (EMD-22814) | #5 Trap1_SdhB$_3$ (EMD-22815) | #6 Trap1_SdhB$_4$ (EMD-22816) (PDB 7KCM) |
|---|---|---|---|---|---|---|
| **Data collection and processing** | | | | | | |
| Microscope | FEI Titan Krios | FEI Titan Krios | FEI Titan Krios | FEI Titan Krios | FEI Titan Krios | FEI Titan Krios |
| Camera | Gatan K2 Summit | Gatan K2 Summit | Gatan K2 Summit | Gatan K2 Summit | Gatan K2 Summit | Gatan K2 Summit |
| Energy Filter | 20 eV slit | 20 eV slit | 20 eV slit | 20 eV slit | 20 eV slit | 20 eV slit |
| Magnification | 165,000x | 165,000x | 165,000x | 165,000x | 165,000x | 165,000x |
| Voltage (kV) | 300 | 300 | 300 | 300 | 300 | 300 |
| Electron exposure (e–/Å$^2$) | 72 | 72 | 72 | 72 | 72 | 72 |
| Defocus range (μm) | -0.7 to -2.1 | -0.7 to -2.1 | -0.7 to -2.1 | -0.7 to -2.1 | -0.7 to -2.1 | -0.7 to -2.1 |
| Pixel size (Å) | 0.814 | 0.814 | 0.814 | 0.814 | 0.814 | 0.814 |
| Symmetry imposed | C1 | C1 | C1 | C1 | C1 | C1 |
| Micrographs (no.) | 10,723 | 10,723 | 10,723 | 10,723 | 10,723 | 10,723 |
| Initial particle images (no.) | 3,415,199 | 3,415,199 | 3,415,199 | 3,415,199 | 3,415,199 | 3,415,199 |
| Final particle images (no.) | 180,584 | 236,804 | 80,304 | 55,565 | 43,148 | 57,787 |
| Map resolution (Å) | 3.26 | 3.14 | 3.38 | 3.57 | 3.52 | 3.43 |
| FSC threshold 0.143 | | | | | | |
| Map resolution range (Å) | 2.5-4.5 | 2.5-4.5 | 2.5-6.5 | 2.5-6.5 | 2.5-6.5 | 2.5-6.5 |
| **Refinement** | | | | | | |
| Initial model used (PDB code) | 6XG6 | 6XG6,1ZOY | | | | 6XG6,1ZOY |
| Model resolution (Å) | 3.5 | 3.4 | | | | 3.5 |
| FSC threshold 0.5 | | | | | | |
| Map sharpening *B* factor (Å$^2$) | -110 | -109 | | | | -80 |
| Model composition | | | | | | |
| Non-hydrogen atoms | 9825 | 10057 | | | | 10841 |
| Protein residues | 1220 | 1249 | | | | 1347 |
| Ligands | 2 | 2 | | | | 3 |
| Ions | 4 | 4 | | | | 4 |
| *B* factors (Å$^2$) | | | | | | |
| Protein | 72.33 | 56.00 | | | | 69.39 |
| Ligand | 39.65 | 40.71 | | | | 62.64 |
| R.m.s. deviations | | | | | | |
| Bond lengths (Å) | 0.003 | 0.002 | | | | 0.002 |
| Bond angles (°) | 0.526 | 0.508 | | | | 0.524 |
| Validation | | | | | | |
| MolProbity score | 1.80 | 1.85 | | | | 1.81 |
| Clashscore | 9.34 | 10.52 | | | | 9.35 |
| Poor rotamers (%) | 0.00 | 0.00 | | | | 0.00 |
| Ramachandran plot | | | | | | |
| Favored (%) | 95.61 | 95.53 | | | | 95.49 |
| Allowed (%) | 4.39 | 4.47 | | | | 4.51 |
| Disallowed (%) | 0.00 | 0.00 | | | | 0.00 |
